# Detection of multiple circulating *Leishmania* species in *Lutzomyia longipalpis* in the city of Governador Valadares, southeastern Brazil

**DOI:** 10.1101/445379

**Authors:** Mariana Santos Cardoso, Gabrielle Ariadine Bento, Laila Viana de Almeida, Joseane Camilla de Castro, João Luís Reis Cunha, Vanessa de Araújo Barbosa, Cristian Ferreira de Souza, Reginaldo Peçanha Brazil, Hugo Oswaldo Valdivia, Daniella Castanheira Bartholomeu

**Author notes:** Current address: U.S. Naval Medical Research Unit No. 6, Bellavista, Callao, Peru. Corresponding author (DCB).

## Abstract

**Background:** Leishmaniasis encompasses a group of diverse clinical diseases caused by protozoan parasites of the *Leishmania* genus. This disease is a major public health problem in the New World affecting people exposed in endemic regions. The city of Governador Valadares (Minas Gerais/Brazil) is a re-emerging area for visceral leishmaniasis, with 191 human cases reported from 2008 to 2017 and a lethality rate of 14.7%. The transmission of the parasite occurs intensely in this region with up to 22% of domestic dogs with positive serology for the visceral form. *Lu. longipalpis* is one of the most abundant sand fly species in this area. Despite this scenario, so far there is no information regarding the circulating *Leishmania* species in the insect vector *Lutzomyia longipalpis* in this focus.

**Methodology/Principal Findings:** We collected 616 female *Lutzomyia longipalpis* sand flies between January and September 2015 in the Vila Parque Ibituruna neighborhood (Governador Valadares/MG), which is located on a transitional area between the sylvatic and urban environments with residences built near a preserved area. After DNA extraction of individual sand flies, the natural *Leishmania* infections in *Lu. longipalpis* were detected by end-point PCR, using primers derived from kDNA sequences, specific for *L. (Leishmania)* or *L. (Viannia)* subgenus. The sensitivity of these PCR reactions was 0.1 pg of DNA for each *Leishmania* subgenus and the total infection rate of 16.2% (100 positive specimens). Species-specific PCR detected the presence of multiple *Leishmania* species in infected *Lu. longipalpis* specimens in Governador Valadares, including *L. amazonensis* (n=3), *L. infantum* (n=28), *L. (Viannia)* spp. (n=20), coinfections with *L. infantum* and *L. (Viannia)* spp. (n=5), and *L. (Leishmania)* spp (n=44).

**Conclusions:** Our results demonstrate that multiple *Leishmania* species circulate in *Lu. longipalpis* in Governador Valadares and reveal a potential increasing risk of transmission of the different circulating parasite species. This information is a key factor for planning surveillance and effective control strategies against leishmaniasis in this endemic focus.

**Author summary:** Leishmaniasis is a neglected tropical disease transmitted to mammals by the bite of sand flies infected with parasites of the *Leishmania* genus. This disease affects millions of people in various regions of the world, including Brazil. The municipality of Governador Valadares (Minas Gerais/Brazil) is a re-emergent focus of intense transmission of leishmaniasis, with a high number of human cases and a high prevalence of infected domestic dogs. To develop better leishmaniasis control strategies for the region, we performed a surveillance study of *Lu. longipalpis,* the main vector of visceral leishmaniasis in Brazil, and identified circulating species of *Leishmania* in this insect vector. We estimate that the natural infection rate of *Lu. longipalpis* for these parasites was of 16.2% in the study area. We also detected the presence of multiple circulating *Leishmania* species *(L. amazonensis, L. infantum* and *Viannia* subgenus) in *Lu. longipalpis* in Governador Valadares city, including 5 sand flies coinfected with *L. infantum* and *L. (Viannia).* Thus, our results reinforce the need for a rigid and systematic control of the sand flies monitoring in this area, due to the potential risk of transmission of different species of the *Leishmania* parasites.

## Introduction

Leishmaniasis is a protozoan parasitic infection transmitted to mammals by the bite of infected phlebotomines (Diptera: Psychodidae). This disease remains as an important illness in the world with an estimate of more than 1 billion people living in endemic areas at risk of infection and 12 million people infected [1]. Leishmaniasis encompasses a wide range of clinical manifestations that are classified into two major types that are the visceral and tegumentary (cutaneous and mucocutaneous) forms. It is reported that visceral leishmaniasis (VL) accounts for 0.2 to 0.4 million cases per year whereas tegumentary leishmaniasis (TL) for 0.7 to 1.2 million [2]. In South America, Brazil is one of the most endemic regions for both VL and TL. Epidemiological data indicates that this country is among the six countries that contribute with 90% of all VL cases in world and among the 10 countries that report between 70 to 75% of all TL cases [2].

The transmission cycle of *Leishmania* parasites relies on the bite of infected sand flies belonging to the genus *Phlebotomus* in the Old World and *Lutzomyia* in the New World. Entomological studies have shown that 70 out of 900 sand fly species (Diptera: Psychodidae: Phlebotominae) are implicated in leishmaniasis transmission [3-5]. For this reason, the identification of vectors and the assessment of natural *Leishmania* infections are essential for epidemiological control and estimation of risk in endemic areas.

In Brazil, the main causative agent of VL is *Leishmania (Leishmania) infantum*, which is mainly transmitted by *Lu. longipalpis* [6]. This sand fly species is spread in most Brazilian states and has shown to be highly adaptable to urban environments [7-10]. The domestic dog *(Canis familiaris)* is considered the main domestic reservoir and a key component of the VL epidemiological chain in urban areas [11, 12].

An important concern about leishmaniasis is the re-emergence of this disease in previously controlled settings and the adaptation into urban environments. These recent events are the result of complex factors including migration, climate change and deforestation [13].

The municipality of Governador Valadares in the southeastern Brazilian state of Minas Gerais is one of these re-emerging areas. This city was an area of intense VL transmission in the 1960s, when a VL control strategy was adopted. This strategy led to a reduction of VL cases up to the beginning of the 1990s when control efforts and epidemiological surveillance were interrupted [11]. VL reappeared again in 2008 when the first human cases were reported [11] and the disease continued to expand reaching up to 191 human VL cases until 2017 [14] and reports of up to 22% of domestic dogs positive by serology in the period 2014-2015 [15]. In addition, this city is also considered endemic for TL with 144 cases reported between 2008 and 2017 [16].

The phlebotomine fauna in Governador Valadares is composed of 12 species belonging to the *Brumptomyia* and *Lutzomyia* genus [11, 17]. Some of these species are known leishmaniasis vectors such as *Lu. intermedia* and *Lu. whitmani*, which could participate in TL transmission in this area [17, 18], and *Lu. longipalpis*, which is one of the most abundant species in peridomestic and domestic areas of the city of Governador Valadares [11, 17]. In other cities of Minas Gerais, several authors have also demonstrated the abundance of *Lu. longipalpis* in urban areas where VL is endemic [7-9].

Recently, our group performed a comparative genomics study using *Leishmania* isolates from dogs with clinical manifestations of VL in Governador Valadares. Molecular genotyping found 34 isolates positive for *L. infantum* and 2 for *L. amazonensis*, an important etiological agent of human cutaneous leishmaniasis [19]. This study evidences the presence and transmission of *L. amazonensis* in southeastern Brazil, an atypical region for this species that was mainly distributed in the north and northeast regions of the country [20, 21].

Herein, we performed a surveillance study of *Lu. longipalpis* in the city of Governador Valadares in order to assess infection rates of sand flies by *Leishmania* and identify circulating species of this parasite in this insect vector. This information will be essential for assessing the risk of transmission and developing better control strategies in this region.

## Methods

### Ethics Statement

This project has authorization and license to capture sand flies under the number 32669-4 IBAMA and with free and informed consent of the owners of the residences.

### Study site and sand fly collections

Sand flies were captured during the year of 2015 in the municipality of Governador Valadares (18° 53’ 5.5” S, 41° 56’ 35.6” W), located in the Doce River Valley of the southeastern Brazilian state of Minas Gerais (Fig 1). This city was chosen primarily due to the high abundance of *Lu. longipalpis* and the emergence of leishmaniasis, being an area considered of intense *L. infantum* transmission [11]. Governador Valadares is an important economic urban center of the Doce River Valley having an estimated population of 280,901 inhabitants [22] and is considered a humid tropical area with average annual temperature and precipitation of 24.2 °C and 1,109 mm, respectively [23].

**Fig 1.**
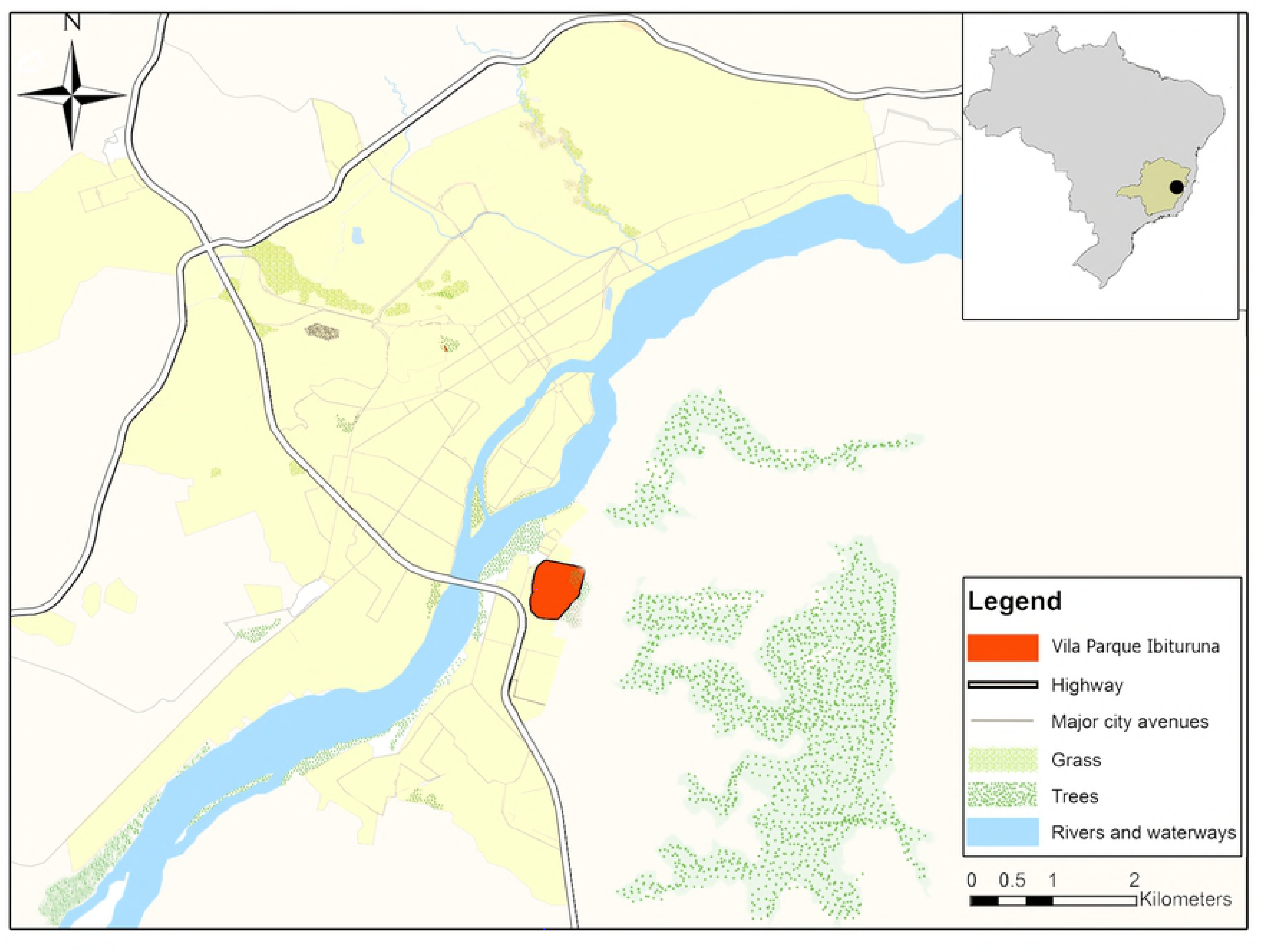
Map of the study area in the city of Governador Valadares. The upper right inset shows the location of the Minas Gerais State (green) in Brazil and the city of Governador Valadares (black circle). On the left, the region marked in orange represents the neighborhood Vila Parque Ibituruna in Governador Valadares, where the sand flies were collected. The map was created using ArcGIS V.10.0.

Sand flies were collected between January and September 2015 in 20 nights of field work activities in the Vila Parque Ibituruna neighborhood (Fig 1). This district is located on a transitional area between rural and urban environments with residences built near the Ibituruna Peak, an environmental preservation area. Two residences were selected for collections based on the presence of gardens, organic matter, chickens and previous *Lu. longipalpis* sampling (unpublished data). At each dwelling, sand flies were collected at the peridomicile using eight HP light traps (4 light traps per garden) equipped with a dispenser that contained the synthetic pheromone (±)-9- methylgermacrene-B to increase capture rates of *Lu. longipalpis* [24].

*Lu. longipalpis* were sexed and the taxonomic identification was based on keys developed by Galati [25, 26]. Male sand flies were confirmed by the presence pale spot on abdominal tergite IV and by the morphological characteristics of the genitalia [25, 26]. *Lu. longipalpis* females were identified according to characteristics of cibarium and spermathecae [25, 26] and were preserved in 6% DMSO for molecular testing.

### DNA extraction and molecular detection of *Leishmania*

Collected *Lu. longipalpis* females and uninfected *Lu. longipalpis* from a colony of the Department of Parasitology of the Federal University of Minas Gerais (UFMG) were submitted to DNA extraction.

DNA was isolated following the protocol previously described by [27] with modifications. Briefly, sand flies were individually placed in 1.5 mL tubes and were homogenized using 50 μl of lysis buffer (0.08 M NaCl, 0.16 M sucrose, 0.06 M EDTA, 0.5% SDS and 0.1 M Tris-HCl, pH 8.6). The homogenate was incubated for 30 min at 65 °C and then 8 M of potassium acetate was added for a final concentration of 1 M and incubated at 4 °C for 30 min. The mixture was subsequently centrifuged at 13,000xg for 10 min at room temperature. The supernatant was transferred to a new tube and mixed with 100 μl of 95% ethanol. After centrifugation at 13,000xg for 10 min, the pellet was washed with 70% ethanol, air dried and resuspended in 50 μl of water.

The presence of *Leishmania* DNA was detected by PCR using subgenus specific primers that target the minicircle kinetoplast DNA (kDNA) of the *Leishmania* or *Viannia* subgenus (Table 1). *Leishmania* subgenus positive samples were subsequently evaluated through species-specific PCR, using primers derived from maxicircle kDNA, which are capable of detecting *L. amazonensis* or *L. infantum* (Table 1).

**Table 1.**
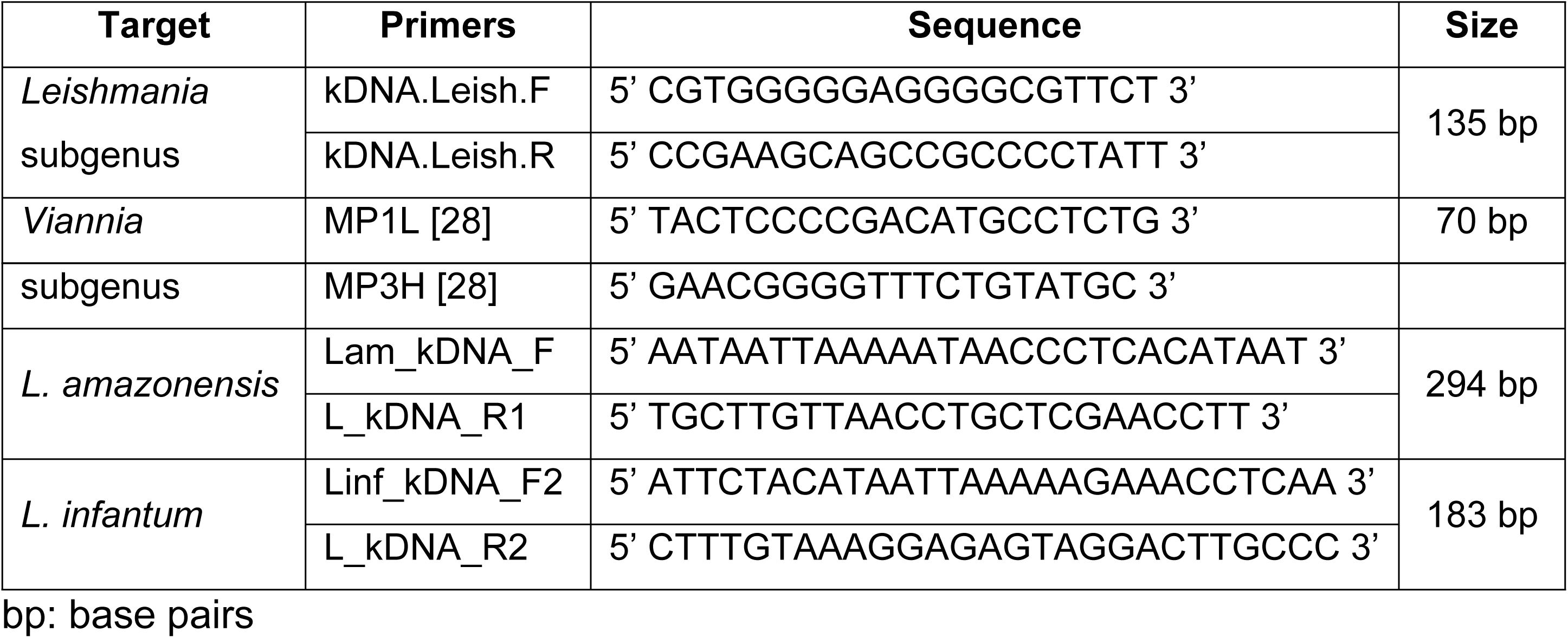
Primers used in PCR reactions with their respective sequences and sizes of amplified products.

The specificity of each pair of primers was checked using 1 ng of the following parasite DNAs: *L. amazonensis* (IFLA/BR/1967/PH8), *L. braziliensis* (MHOM/BR/75/M2904), *L. donovani* (LD1S/MHOM/SD/00-strain1S), *L. guyanensis* (MHOM/BR/1975/M4147), *L. infantum* (MHOM/BR/1974/PP75), *L. lainsoni* (MHOM/BR/1981/M6426), *L. mexicana* (MHOM/BZ/1982/BEL21), *L. naiffi* (MDAS/BR/1979/M5533), *L panamensis* (MHOM/PA/71/LS94), and *L. shawi* (MHOM/BR/96/M15789), under the reaction conditions described below. The sensitivity of PCR was also performed for each pair of primers from serial dilutions of 10 ng to 1 fg of the following DNAs: *L. amazonensis* and *L. infantum* for the primers kDNA.Leish_F/kDNA.Leish_R; *L. braziliensis* for MP1L/MP3H; *L. amazonensis* for Lam_kDNA_F/L_kDNA_R1; and *L. infantum* para Linf_kDNA_F2/L_kDNA_R2.

PCRs were carried out in 20 μL reactions, containing 1X GoTaq Green buffer (Promega), 0.2 mM dNTPs, 0.5 μM of each primer, 1 U of Taq DNA polymerase (Phoneutria), and 1 or 5 μL of DNA. PCRs of sand fly specimens were performed in two steps. Initially, samples were analyzed in pools of 5 sand flies for initial screening with the *Leishmania* and *Viannia* subgenus specific primers. In this case, 5 μL of each pool was used per PCR. Subsequently, individual DNA samples of sand flies of each positive pool were submitted to PCR using the subgenus and species-specific primers. The thermal cycling conditions consisted of an initial denaturation at 94 °C for 5 min; followed by 30 cycles of denaturation at 94 °C for 30 sec, annealing at 56 °C for *Leishmania* subgenus, 60 °C for *Viannia* subgenus and 50 °C for *L. amazonensis* and *L. infantum* for 30 sec, and extension at 72 °C for 30 sec; and a final extension step at 72 °C for 7 min. The expected size of the bands for each reaction is indicated in Table 1. The amplified products were analyzed in 2.0% agarose gel electrophoresis in 1x TAE buffer containing 0.5 μg/mL ethidium bromide and visualized in UV light, using the ImageQuant LAS 4000 (GE Healthcare Life Science).

## Results

### Multiple *Leishmania* species are detected in *Lu. longipalpis* in Governador Valadares

A total of 616 females of *Lu. longipalpis* collected in Governador Valadares were used for molecular detection of *Leishmania* by PCR. The specificity of each primer pair used in this study was assessed in PCR reactions using as template gDNA from 10 different reference *Leishmania* species: 4 *L. (Leishmania)* and 6 *L. (Viannia)* (Table 2 and Fig 2). The primers kDNA.Leish.F/R and MP1L/MP3H [28] were specific for the *Leishmania* and *Viannia* subgenus, respectively. The primers Lam_kDNA_F/L_kDNA_R1 were specific for *L. amazonensis* and *L. mexicana* (both of the *Leishmania mexicana* complex), whereas the primers Linf_kDNA_F2/L_kDNA_R2 presented specificity for *L. donovani* and *L. infantum* (both of the *Leishmania donovani* complex).

**Table 2.** Sensitivity of PCR and specificity of each primer pair used for *Leishmania* detection.

| Primers | Sensitivity | Specificity |
| --- | --- | --- |
| **kDNA *L. (Leishmania)***<br>(kDNA.Leish_F/kDNA.Leish_R) | 100 fg | *Leishmania* subgenus |
| **kDNA *L. (Viannia)***<br>(MP1L/MP3H) | 100 fg | *Viannia* subgenus |
| **kDNA *L. amazonensis***<br>(Lam_kDNA_F/L_kDNA_R1) | 1 pg | *L. amazonensis* and *L. mexicana* |
| **kDNA *L. infantum***<br>(Linf_kDNA_F2/L_kDNA_R2) | 1 pg | *L. infantum* and *L. donovani* |

**Fig 2.**
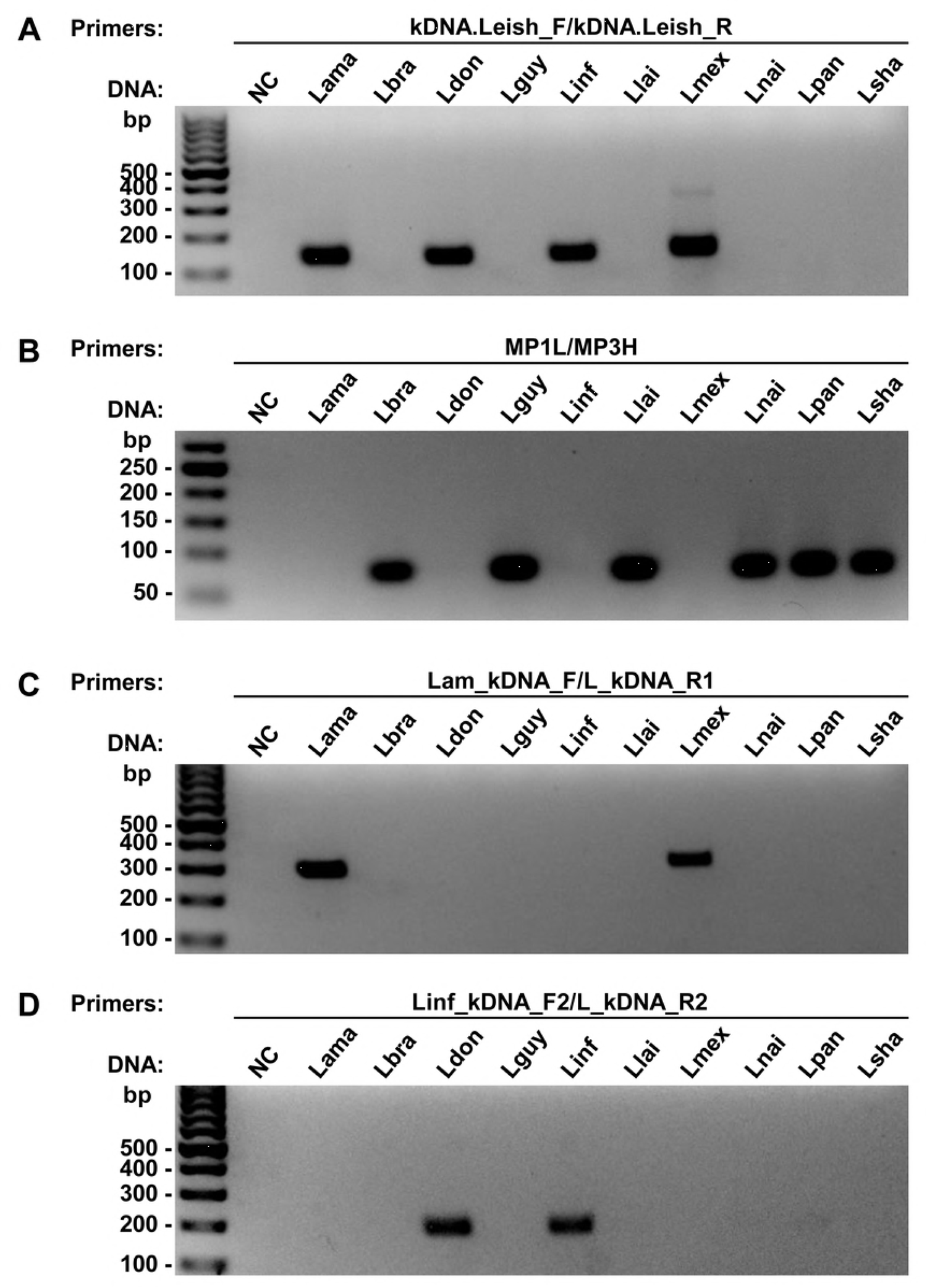
Specificity analysis of each pair of primers using different *Leishmania* species in PCR. **(A)** The pair of primer kDNA.Leish was specific for the *Leishmania* subgenus amplifying the DNA of *L. amazonenis, L. donovani, L. infantum,* and *L. mexicana;* **(B)** primers MP1L/MP3H were specific for the *Viannia* subgenus, amplifying the DNA of *L. braziliensis, L. guyanensis, L. lainsoni, L. naiffi, L. panamensis,* and *L. shawi;* **(C)** the primers Lam_kDNA_F/L_kDNA_R1 amplify both *L. amazonenis* and *L. mexicana;* and **(D)** the primers Linf_kDNA_F2/L_kDNA_R2 amplify *L. donovani* and *L. infantum.* bp: molecular size marker in base pairs; NC: negative control (without DNA); DNA: Lama: *L. amazonensis;* Lbra: *L. braziliensis;* Ldon: *L. donovani;* Lguy: *L. guyanensis;* Linf: *L. infantum;* Llai: *L. lainsoni;* Lmex: *L. mexicana;* Lnai: *L. naiffi;* Lpan: *L. panamensis;* Lsha: *L. shawi*.

The gDNAs of *L. amazonensis* and *L. infantum* for the primers kDNA.Leish_F/kDNA.Leish_R, *L. braziliensis* for MP1L/MP3H, *L. amazonensis* for Lam_kDNA_F/L_kDNA_R1, and *L. infantum* for Linf_kDNA_F2/L_kDNA_R2 were diluted from 10 ng to 1 fg to evaluate the sensitivity of PCR. The *Leishmania* species used to check the specificity of each primer were *L. amazonensis, L. braziliensis, L. donovani, L. guyanensis, L. infantum, L. lainsoni, L. mexicana, L. naiffi, L. panamensis,* and *L. shawi.* gDNA: genomic DNA.

The limit of detection for each PCR was also evaluated using serial dilutions of gDNA (10 ng to 1 fg) from *L. amazonensis, L. braziliensis,* and *L. infantum* (Table 2 and S1 Fig). The *Leishmania* and *Viannia* subgenus specific PCR, using primers derived from the minicircle kDNA, had a limit of detection of 100 fg of genomic DNA of the parasite. On the other hand, the PCR using primers derived from maxicircules kDNA of the *L. amazonensis* and *L. infantum* species presented a limit of detection of 1 pg.

To identify sand flies that were infected with parasites of the *Leishmania* and *Viannia* subgenus, we prepared a pool of gDNA containing samples of 5 sand flies and tested by end-point PCR, using primers kDNA.Leish.F/R or MP1L/MP3H, respectively (S2 and S3 Figs). Then DNA samples from positive pools for each subgenus were individually tested with these same primers derived from kDNA of *Leishmania* (Fig 3A) and *Viannia* subgenus (Fig 3B). Gels with all samples tested individually are in S4 and S5 Figs. We identified 80 positive samples of sand flies for *Leishmania* subgenus and 25 for *Viannia* subgenus.

**Fig 3.**
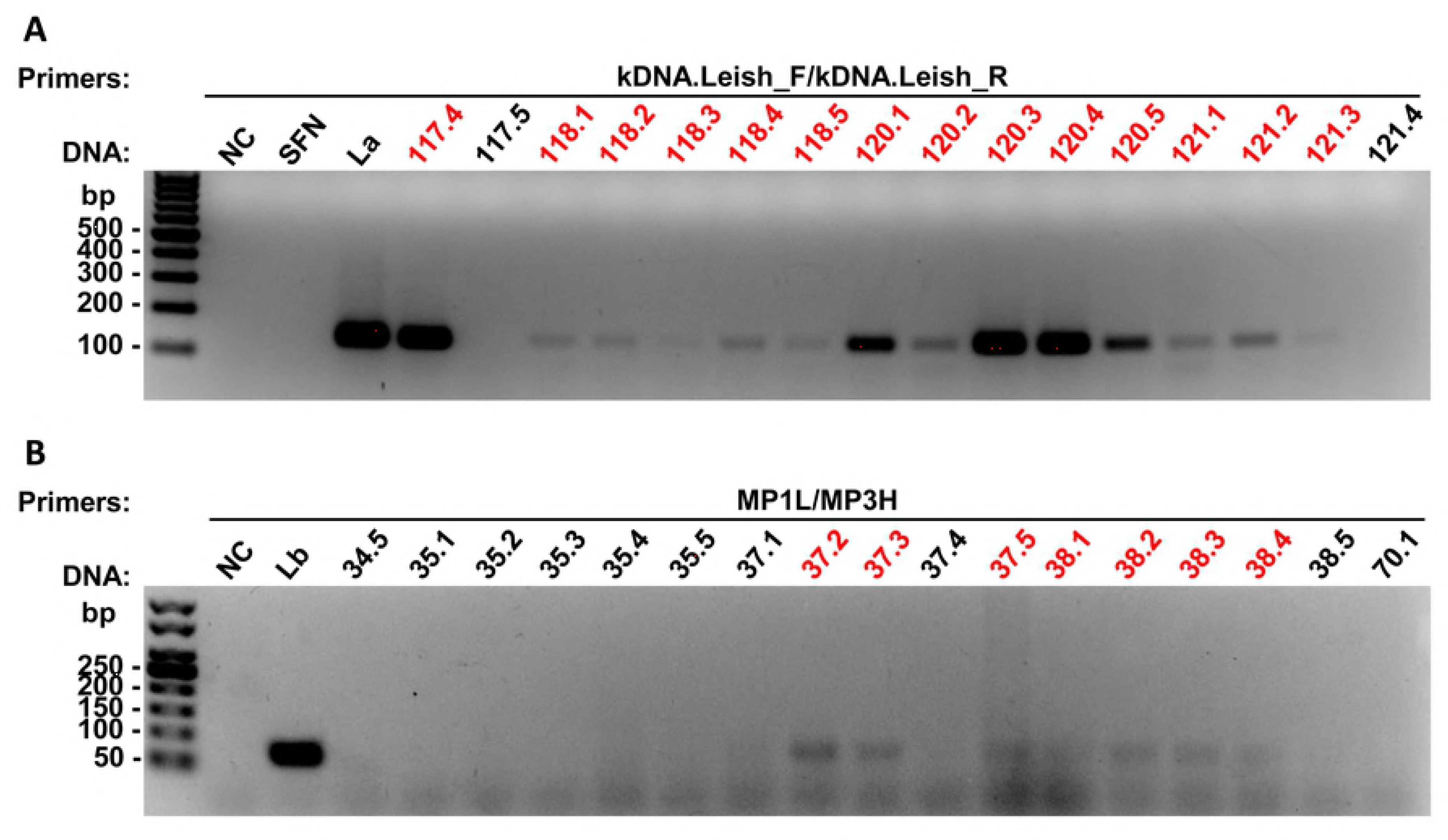
PCR analysis of *Lu. longipalpis* gDNA, using primers derived from *Leishmania* and *Viannia* subgenus kDNA. Representative gels of the PCR with individual DNA samples of sand flies, using the primers kDNA.Leish, which amplify *Leishmania* subgenus with a fragment of 135 bp **(A)**, and MP1L/M3H, which amplify *Viannia* subgenus, with a fragment of 70 bp **(B)**. Samples marked in red were positive for the respective primers tested. bp: molecular size marker in base pairs; NC: negative control (without DNA); SFN: not infected sand fly; DNA control: La: *L. amazonensis;* Lb: *L. braziliensis*.

Of the 80 positive samples for the *Leishmania* subgenus, 3 were positive for *L. amazonensis* (Fig 4A and S6 Fig) and 33 for *L. infantum* (Fig 4B and S7 Fig). We were unable to identify the species from the remaining 44 samples due to the lower sensitivity of the species-specific primers. Additionally, we found 5 sand flies potentially coinfected with *L. infantum* and a member of the *Viannia* subgenus (Fig 5).

**Fig 4.**
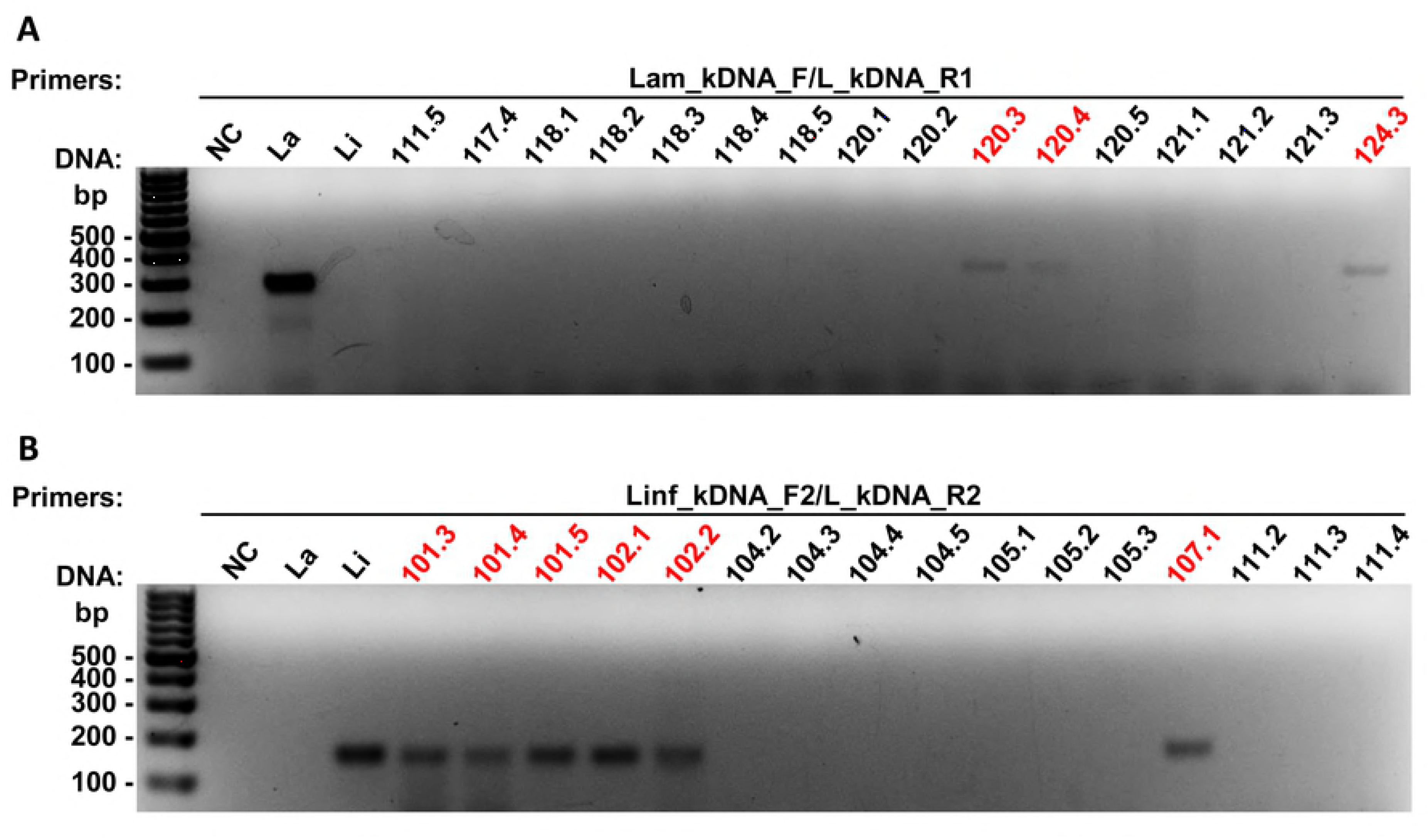
PCR analysis of *Lu. longipalpis* gDNA, using primers derived from *L. amazonensis* and *L. infantum* kDNA. Representative gels of the PCR with individual DNA samples of sand flies, using the primers Lam_kDNA_F/L_kDNA_R1, which amplify *L. amazonensis* with a fragment of 294 bp **(A)**, and Linf_kDNA_F2/L_kDNA_R2, which amplify *L. infantum* with a fragment of 183 bp **(B)**. Samples marked in red were positive for the respective primers tested. bp: molecular size marker in base pairs; NC: negative control (without DNA); SFN: not infected sand fly; DNA control: La: *L. amazonensis;* Li: *L. infantum*.

**Fig 5.**
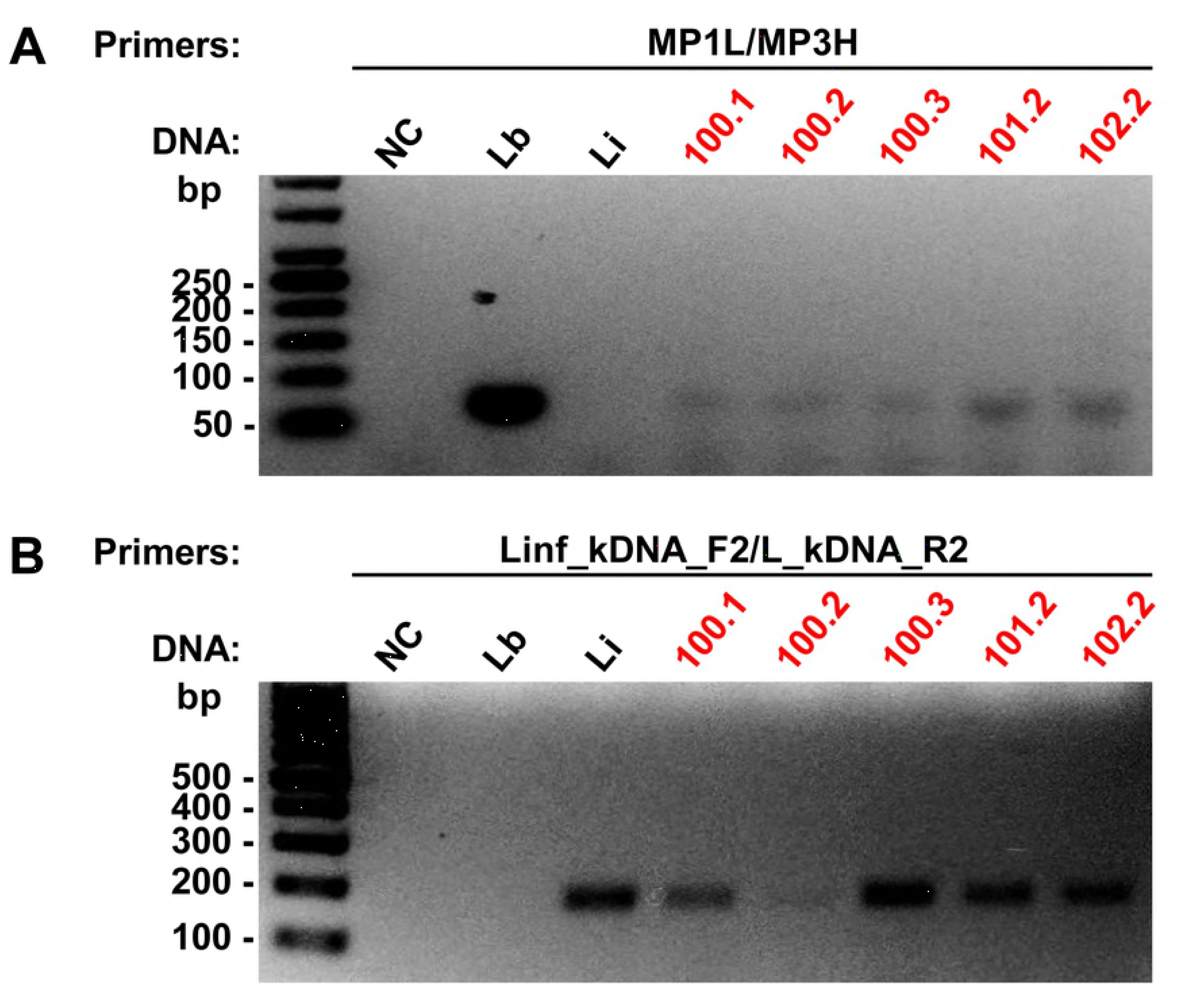
PCR analysis of *Lu. longipalpis* gDNA coinfected with *L. infantum* and *L. (Viannia)* spp. Gels of the PCR with the positive samples both for *Viannia* subgenus, using the primers MP1L/MP3H (fragment: 70 bp) **(A)**, and for *L. infantum,* using the primers Linf_kDNA_F2/L_kDNA_R2 (fragment: 183 bp) **(B**. Samples marked in red were positive for the respective primers tested. bp: molecular size marker in base pairs; NC: negative control (without DNA); SFN: not infected sand fly; DNA control: Lb: *L. braziliensis;* Li: *L. infantum*.

The estimated natural infection rate of *Lu. longipalpis* for the *Leishmania* genus was of 16.2%. The natural infection rates for the *Leishmania* and *Viannia* subgenus were of 13% and 4%, respectively (Fig 6).

**Fig 6.**
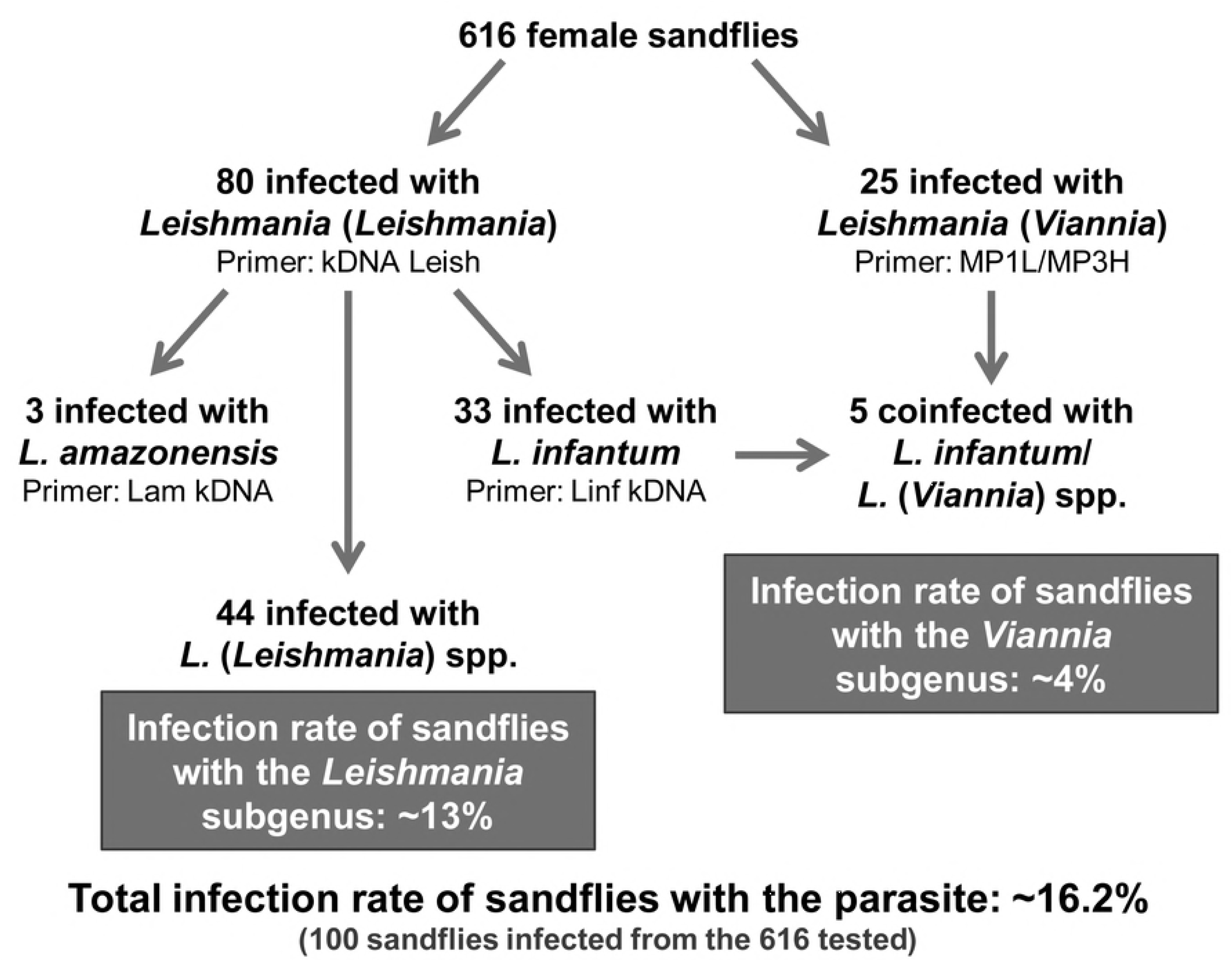
Overview of the results obtained in the study of the *Lu. longipalpis* infection with *Leishmania*. The number of infected sand flies and the total and subgenus infection rate are shown in the diagram.

## Discussion

Governador Valadares is a re-emergent focus of intense transmission of TL and VL with a high number of human cases and a high prevalence of infected domestic dogs. After the interruption of the control program in the early 1990s, there was a reemergence of cases in the region with a lethality rate of more than 16% [11, 17]. In a study conducted in 2013, the average prevalence of positive dogs was 30.2%, reaching 53% in some neighborhoods [11]. In another study conducted in 2014 and 2015, seroprevalence rate in dogs tested by DPP was 34.8%, of which 22% were confirmed by ELISA [15]. In addition to VL cases caused by *L. infantum*, our group presented the first report of *L. amazonensis* in Governador Valadares isolated from bone marrow and lymph node aspirates of dogs with visceral symptomatology [19]. Control activities should therefore consider the presence of *L. amazonensis* in this endemic visceral leishmaniasis site and address putative vectors and the risks of coinfections in humans.

In this endemic focus, there were reports of more than 12 sand fly species circulating in the peridomicile areas, with *Lu. longipalpis* being one of the most abundant sand fly species [11, 17]. Despite this scenario, to our knowledge, no study was conducted in the area to investigate the species of *Leishmania* that circulate in this vector. Therefore, in order to investigate the circulating *Leishmania* species in *Lu. longipalpis*, the main vector of VL in Brazil [6], we collected sand flies in Governador Valadares city during the year 2015 and the presence of this parasite was molecularly analyzed by PCR.

The entomological survey was performed in the neighborhood of Vila Parque Ibituruna, where phlebotomines were captured in residences built near the preserved area and having gardens, chicken coop and organic matter. Sand flies are typically associated with sites that exhibit shade, decomposing organic matter, vegetation, and moisture [29].

The occurrence of naturally infected sand flies with *Leishmania* is an important evidence to investigate its role as a vector. However, a number of criteria have to be considered to a phebotomine be incriminated as a natural vector [30, 31]. Two approaches have been used to identify the presence of *Leishmania* DNA in sand flies. The gold-standard method used to study the rate of natural infection in endemic areas has been the digestive tract dissection of female sand flies, permitting the direct observation of the parasites [32]. However, this technique is laborious, time-consuming, difficult to process a large number of samples and does not allow the identification of genus and species [32, 33]. Alternatively, molecular techniques, such as PCR, are highly specific and more sensitive, allowing the detection of a single parasite depending on the used primer [34, 35]. The use of PCR for detection of *Leishmania* DNA in sand flies is a useful technique for the identification of putative vectors in different geographical areas [33, 36-39]. Some targets used to analyze the infection in sand flies are ITS1 [33, 37, 38] and kDNA [36, 40, 41].

In this study, we used primers derived from the minicircle kDNA of the *Leishmania* and *Viannia* subgenus [28], which were highly specific for each evaluated subgenus. We also used primers derived from the maxicircle kDNA of *L. amazonensis* and *L. infantum,* which showed high specificity, although they also recognize another species of the same taxonomic complex. However, the species *L. mexicana* and *L. donovani*, which are also detected by the respective primers, are not found naturally in Brazil.

Primers derived from kDNA are suitable molecular markers for *Leishmania* detection in sand flies, because they are based on sequences with a high copy number per cell, which confers a high sensitivity to the technique [41, 42]. We obtained PCR with sensitivity of 0.1 pg for specific subgenus reactions, and 1 pg for the reactions that amplify *L. amazonensis* and *L. infantum* kDNA sequences. These sensitivities were similar to those obtained in other studies using ITS1 primers [35, 37].

Due to the high prevalence of *Lu. longipalpis* in the municipality of Governador Valadares, we investigated the circulating *Leishmania* species in this vector through molecular detection by PCR. We identified 80 positive sand flies for *Leishmania* subgenus and 25 for *Viannia* subgenus. Of the positive samples for *Leishmania* subgenus, 3 presented DNA of *L. amazonensis*, 33 of *L. infantum*, and 44 remaining without species identification, because of the lower sensitivity of species-specific primers. Interestingly, we also observed 5 sand flies presenting DNA of *L. infantum* and of a representative of *Viannia* subgenus.

*Lu. longipalpis* is the main vector of *L. infantum* in Brazil [6], however, reports of PCR detection of other *Leishmania* species in this sand flies have been described in several studies [33, 38, 43, 44]. In agreement with our results, these studies described the association of *Lu. longipalpis* with *L. infantum,* and also with *L. amazonensis* [43] and *L. braziliensis* (or a parasite belonging to *Viannia* subgenus) [33, 38, 43, 44]. Mixed infections of *L. (Leishmania)* sp. and *L. (Viannia)* sp. were also found in 0.5% of *Lu. longipalpis* evaluated by [43]. However, these findings are not sufficient to incriminate *Lu. longipalpis* as a vector of other species of *Leishmania.*

Studies on experimental infections in sand flies with different *Leishmania* species suggest that *Lu. longipalpis* is a permissive vector, which support the development of different *Leishmania* species [45-47]. The susceptibility of *Lu. longipalpis* to *L. amazonensis, L. braziliensis, L. guyanensis, L. infantum* and *L. mexicana* were experimentally studied by [45]. Only 9% of blood fed sand flies on the lesions of hamsters infected presented the parasite *L. braziliensis* or *L. mexicana* after dissection. A higher infection rate was observed after exposure of the sand fly to *L. amazonensis* (37%) and *L. guyanensis* (100%). Similar results were observed by [47], with two isolates of *L. amazonensis* presenting more than 60% of infection in *Lu. longipalpis* and one isolate of *L. braziliensis* presenting only 5% of infection rate. However, additional studies are necessary to define the role of this sand fly species in the epidemiological context of leishmaniasis and to assess the vectorial competency of *Lu. longipalpis* as putative vector of other *Leishmania* species.

Our results indicate that the natural infection rate of *Lu. longipalpis* with the parasites of *Leishmania* genus was 16.2%, being 13% with *Leishmania* subgenus and 4% with *Viannia* subgenus. These rates are higher than those found in other regions of Brazil [33, 38, 43, 44, 48], which may contribute to the high risk of *Leishmania* infection in Governador Valadares. In studies carried out in Brazil, the natural infection rate of *Lu. longipalpis* was 2.76% for *Leishmania* spp. (0.25% *L. infantum,* 0.75% *L. (Viannia)* sp., 1.25% *L. amazonensis,* and 0.50% mixed infections of *L. (Leishmania)* sp. and *L. (Viannia)* sp. [43], 0.94% for *L. braziliensis* [44], 1.44% for *Leishmania* spp. (0.72% for *L. (Viannia)* sp. and *L. infantum)* [33], 2.79% for *L. infantum* and *L. braziliensis* (1.01% and 1.77%, respectively) [38], and 6.91% for *L. infantum, L. braziliensis* and *L. lainsoni* (3.72%, 2.13% and 1.06%, respectively) [48].

Therefore, this study demonstrates the urgent need for constant surveillance and control of leishmaniasis in the municipality of Governador Valadares, by sand fly population monitoring and seropositive dogs. Further studies are needed to incriminate *Lu. longipalpis* as vector of multiple *Leishmania* species in this endemic focus.

## Disclaimer

HOV is an employee of the US Government. The views expressed in this article are those of the authors and do not necessarily reflect the official policy or position of the Department of the Navy, Department of Defense, nor the US Government.

## Supporting information

**S1 Fig. Sensitivity analysis of PCR for each pair of primers from serial dilutions of *Leishmania* gDNA**. The detection limit of each PCR is indicated by the red arrow: **(A)** 100 fg of *L. amazonensis* and *L. infantum* gDNA for PCR with pair of primer kDNA.Leish; **(B)** 100 fg of *L. braziliensis* gDNA for primers MP1L/MP3H; **(C)** 1 pg of *L. amazonensis* gDNA for primers Lam_kDNA_F/L_kDNA_R1; and **(D)** 1 pg of *L. infantum* gDNA for primers Linf_kDNA_F2/L_kDNA_R2. bp: molecular size marker in base pairs; NC: negative control (without DNA); gDNA: genomic DNA.

**S2 Fig. PCR analysis of pool gDNA of *Lu. longipalpis*, using primers derived from *Leishmania* subgenus kDNA**. Gels of the PCR with pool gDNA sand flies, using the pair of primers kDNA.Leish, which amplifies *Leishmania* subgenus with a fragment of 135 bp. Each pool contains 5 DNA samples of *Lu. longipalpis,* totaling 616 sand flies distributed in 124 pools. Pools marked in red were positive for the primers tested. bp: molecular size marker in base pairs; NC: negative control (without DNA); SFN: not infected sand fly; DNA control: La: *L. amazonensis.*

**S3 Fig. PCR analysis of pool gDNA of *Lu. longipalpis*, using primers derived from *Viannia* subgenus kDNA**. Gels of the PCR with pool gDNA sand flies, using the pair of primers MP1L/MP3H, which amplifies *Viannia* subgenus with a fragment of 70 bp. Each pool contains 5 DNA samples of *Lu. longipalpis,* totaling 616 sand flies distributed in 124 pools. Pools marked in red were positive for the primers tested. bp: molecular size marker in base pairs; NC: negative control (without DNA); SFN: not infected sand fly; DNA control: Lb: *L. braziliensis.*

**S4 Fig. PCR analysis of individual gDNA of *Lu. longipalpis*, using primers derived from *Leishmania* subgenus kDNA.** Gels of the PCR with individual DNA samples from positive pools of sand flies, using the pair of primers kDNA.Leish, which amplifies *Leishmania* subgenus with a fragment of 135 bp. We detected 80 sand flies infected with *L. (Leishmania)* spp. Samples marked in red were positive for the primers tested. bp: molecular size marker in base pairs; NC: negative control (without DNA); SFN: not infected sand fly; DNA control: La: *L. amazonensis.*

**S5 Fig. PCR analysis of individual gDNA of *Lu. longipalpis*, using primers derived from *Viannia* subgenus kDNA**. Gels of the PCR with individual DNA samples from positive pools of sand flies, using the pair of primers MP1L/MP3H, which amplifies *Viannia* subgenus with a fragment of 70 bp. We detected 25 sand flies infected with *L. (Viannia)* spp. Samples marked in red were positive for the primers tested. bp: molecular size marker in base pairs; NC: negative control (without DNA); SFN: not infected sand fly; DNA control: Lb: *L. braziliensis.*

**S6 Fig. PCR analysis of individual gDNA of *Lu. longipalpis*, using primers derived from *L. amazonensis* kDNA**. Gels of the PCR with positive DNA samples for *Leishmania* subgenus, using the pair of primers Lam_kDNA_F/L_kDNA_R1, which amplifies *L. amazonensis* with a fragment of 294 bp. We detected 3 sand flies infected with *L.* amazonensis. Samples marked in red were positive for the primers tested. bp: molecular size marker in base pairs; NC: negative control (without DNA); SFN: not infected sand fly; DNA control: La: *L. amazonensis;* Li: *L. infantum.*

**S7 Fig. PCR analysis of individual gDNA of *Lu. longipalpis*, using primers derived from *L. infantum* kDNA**. Gels of the PCR with positive DNA samples for *Leishmania* subgenus, using the pair of primers Linf_kDNA_F2/L_kDNA_R2, which amplifies *L. infantum* with a fragment of 183 bp. We detected 33 sand flies infected with *L.* infantum. Samples marked in red were positive for the primers tested. bp: molecular size marker in base pairs; NC: negative control (without DNA); SFN: not infected sand fly; DNA control: La: *L. amazonensis;* Li: *L. infantum.*

